# Transplanted pluripotent stem cell-derived photoreceptor precursors elicit conventional and unusual light responses in mice with advanced retinal degeneration

**DOI:** 10.1101/2020.09.22.308726

**Authors:** Darin Zerti, Gerrit Hilgen, Birthe Dorgau, Joseph Collin, Marius Ader, Lyle Armstrong, Evelyne Sernagor, Majlinda Lako

## Abstract

Retinal dystrophies often lead to blindness. Developing therapeutic interventions to restore vision is therefore of paramount importance. Here we demonstrate the ability of pluripotent stem cell-derived cone precursors to engraft and restore light responses in the *Pde6brd1* mouse, an end-stage photoreceptor degeneration model. Up to 1.5% of precursors integrated into the host retina, differentiated into cones and formed synapses with bipolar cells. Half of the transplanted mice exhibited visual behaviour and 33% showed binocular light sensitivity. The majority of ganglion cells exhibited contrast-sensitive ON, OFF or ON-OFF light responses and even motion sensitivity. Many cells also exhibited unusual responses (e.g. light-induced suppression), presumably reflecting remodelling of the neural retina. Our data indicate that despite relatively low engraftment yield, engrafted pluripotent stem cell-derived cone precursors can elicit light responsiveness even at advanced degeneration stages. Further work is needed to improve engraftment yield and counteract retinal remodelling to achieve useful clinical applications.

## INTRODUCTION

Characterized by dysfunction and ensuing progressive cellular loss, retinal degenerative diseases (RDDs) are the leading cause of blindness and currently lack effective treatment. In particular, vision impairment due to photoreceptor degeneration affects more than 43 million people globally, with an exponential increase in the ageing population predominantly over 50 years. Mammalian retinal degeneration initiated by gene defects in rods, cones or the retinal pigmented epithelium (RPE) almost unilaterally trigger the loss of the sensory retina, effectively leaving the neural retina deafferented. The latter responds to this challenge by remodelling surviving cells and connectivity (Marc et al., 2003). All RDDs have the same outcome: photoreceptor loss followed by sustained cell death and remodelling, leading to profound morphological, biochemical and physiological alterations that involve all structures, including neurons, blood vessels and glial cells (Marc et al., 2003). These changes have been extensively described in animal models such as *Pde6brd1* mice, an animal model of autosomal recessive form of Retinitis Pigmentosa (RP) (Farber et al., 1994; L D Carter-Dawson; et al., 1978). Retinal remodelling occurs in 3 phases (Jones and Marc, 2005): 1) primary photoreceptor loss; 2) secondary photoreceptor degeneration, microglial proliferation, Müller glia activation leading to seal formation between the retina and the choroid; 3) de *novo* neurite formation, rewiring and neuronal death, posing an enormous challenge to successful potential therapies based on genetic, cellular and bionic approaches.

Transplantation of photoreceptor progenitors in the degenerating retina is a promising approach, but in order to be successful, it requires that the transplanted cells engraft efficiently and establish functional synaptic connections with the host tissue (more specifically, with bipolar and horizontal cells), which would be reflected by de *novo* responses to light in bipolar and retinal ganglion cells (RGCs). One of the major obstacles resulting from remodelling of the neural retina is the development of pathological strong, continuous oscillations in the inner retina (Menzler and Zeck, 2011; Stasheff, 2008). These oscillations occur at a frequency of about 10 Hz, dominating activity in the RGC layer in the form of slow local field potentials and rhythmic bursting. These oscillations arise from cross-junctional interactions between AII amacrine/ON cone bipolar cells (Choi et al., 2014; Trenholm et al., 2012). Because of the very poor signal-to-noise ratio triggered by these oscillations, potential rescued light responses originating from transplanted photoreceptors are very unlikely to be detected at the RGC level, and even less so in retinal central projections. It has been shown by our group (Barrett et al., 2015, 2016) and others (Toychiev et al., 2013) that gap junction blockade is a very useful approach to dampen these oscillations, thereby significantly improving the signal-to-noise ratio of responses to light in RGCs.

Replacement of the lost photoreceptors by transplantation offers promising prospects for the reversal of end-stage degeneration (Gonzalez-Cordero et al., 2017), and this therapeutic approach has been widely investigated over the last three decades. Since the introduction of human embryonic stem cells (hESCs) (Thomson et al., 1998) and induced pluripotent stem cells (hiPSCs) (Takahashi et al., 2007), several approaches have been investigated for the generation of retinal cells in vitro. Seminal work from the Sasai lab demonstrated the feasibility of generating murine 3D retinal organoids that closely follow *in vivo* retinogenesis (Eiraku et al., 2011); they subsequently provided a protocol for the generation of human 3D retinal tissue from hESCs with a typical laminated retinal structure, harbouring high numbers of photoreceptors and all other key retinal cell types (Nakano et al., 2012). Since then, several groups, including ours, have optimised differentiation techniques (Boucherie et al., 2011; Capowski et al., 2019; Dorgau et al., 2019; Gonzalez-Cordero et al., 2013; Mellough et al., 2015, 2019; Phillips et al., 2018; Zhong et al., 2014) and shown that these *in vitro* derived retinal organoids respond to light, exhibiting immature responses that are similar to those seen in neonatal mice (Hallam et al., 2018). Because of their unlimited ability to proliferate and intrinsic ability to generate light responsive laminated retinal organoids, both hESCs and hiPSCs represent the most attractive cell source for generating transplantable photoreceptors.

To date, there have been several animal studies evaluating the transplantation of photoreceptor precursors, derived both from human or mouse iPSCs and ESCs, with a few reporting improvements in visual function (Barnea-Cramer et al., 2016; Lamba et al., 2009; Pearson et al., 2012; Singh et al., 2013). Using fluorescent reporters to track the fate of donor-transplanted cells, these studies have shown photoreceptor integration into the outer nuclear layer (ONL), the development of outer and inner segments and typical photoreceptor morphology, as well as the ability to form synapses with host cells. Until recently, it had been assumed that the reported improvements in visual function, at least in those models where some endogenous photoreceptors survived, were due to donor cell integration within the host retina. However, with the recent and unexpected identification of cytoplasmic material exchange from donor to host photoreceptors, most of the former findings regarding donor cell migration, integration in the host retinal circuitry with correct synaptic contacts need to be revisited, especially in animal models where the host photoreceptors survive to some extent (Nickerson et al., 2018; Pearson et al., 2016; Santos-Ferreira et al., 2016).

In 2016, our group reported the successful generation of a reporter labelled hESC line, in which expression of the green fluorescent protein (GFP) is controlled by CRX, a key transcription factor in retinal development with predominant expression in post-mitotic photoreceptor precursors (Collin et al., 2016). We subsequently demonstrated the successful engraftment capability of these photoreceptor precursors in the degenerative retina of *Pde6brd1* mice (Collin et al., 2019). Using a combination of methods, such as the presence of species-specific antigens, immunostaining for human cytoplasmic components and human nuclear antigens and the host-donor nuclei measurement, we were able to exclude cellular transfer between donor and host photoreceptors and demonstrate genuine engraftment of CRX^+^ cone photoreceptors in the putative ONL of *Pde6brd1* mice. Here, we investigate the ability of transplanted photoreceptors to restore light-responses in RGCs in *Pde6brd1* mice. By dampening the spontaneous oscillations in RGCs using gap junction blockade and screening the responses to light (contrast steps and moving gratings) in hundreds to thousands of RGCs at near pan retinal level with a next generation large-scale high-density multielectrode array (MEA), we demonstrate that this approach successfully restores responsiveness to light in RGCs in retinas with end-stage degeneration.

## RESULTS

### Recovery of basic visual function upon transplantation of CRX^+^ human photoreceptors in mice with end-stage retinal degeneration

Photoreceptors are terminally differentiated sensory cells, which once lost, cannot regenerate. In earlier studies, we have shown that hESC-derived CRX+ photoreceptor precursors have the ability to engraft into *Pde6brd1* mice, mature into cones (Collin et al., 2019) and establish contacts with rod bipolar axon terminals. In this study, we assessed whether these transplanted photoreceptors can restore to some extent visual function. At day 90 of differentiation, retinal organoids were collected and dissociated to single cells. CRX-GFP^+^ cells were isolated by fluorescence-activated cell sorting and transplanted bilaterally into the subretinal space of 28 immunosuppressed 10-12 weeks old *Pde6brd1* mice as described previously (Collin et al., 2019). Five additional animals were injected with the vehicle solution (HBSS) (sham group). All injected eyes were examined 3-4 weeks after grafting. GFP^+^ cells were found next to the host inner nuclear layer (INL) (815 ±298 cells per retina, **Fig. 1A**) and these were localised mainly around the injection site (**Fig. 1B**). We confirmed the photoreceptor identify of human donor cells by co-expression of GFP with the pan-photoreceptor marker Recoverin in the putative host ONL in transplanted retinal wholemounts (**Fig. 1C**). No GFP^+^ cells were detected in the sham treated group (data not shown).

**Fig. 1.**
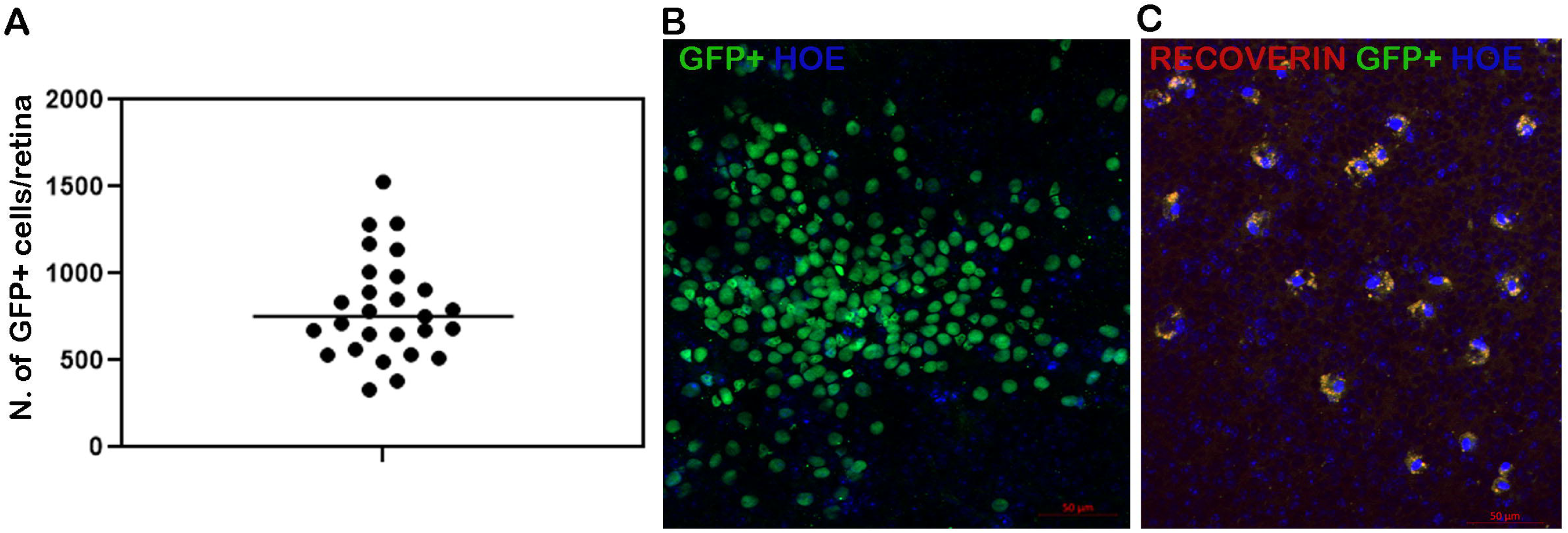
Engraftment of CRX-GFP+ cells into *Pde6brd1* retinas. **(A)** Scatter plot showing the number of GFP^+^ cells per retina per mouse. Data are presented as number of GFP^+^ cells ± STD; (N= 27 retinas); (**B**) Whole mount view of GFP^+^ cells in *Pde6brd1* mice retinas transplanted with CRX^+^ cells; (**C**) Immunohistochemical analysis for Recoverin expression in these transplanted retinas. Scale bars, 50 µm (B, C). Abbreviations: GFP, green fluorescent protein, Hoe, Hoechst. Note: GFP in (B) and (C) represents endogenous GFP expression.

In the mammalian retina, bipolar cells receive signals from photoreceptors and send integrated signals to RGCs. Photoreceptor cell axon terminals form synapses with bipolar cell dendrites and horizontal cell processes in the outer plexiform layer (OPL), and bipolar cell axon terminals connect with amacrine cell and ganglion cell dendrites in the inner plexiform layer (IPL). To investigate synapse formation between the donor and host, we stained the transplanted retinas with a G-protein subunit G0α antibody, a marker for rod and cone ON bipolar cells dendrites (Haverkamp and Wässle, 2000; Hilgen et al., 2012; Vardi, 1998). G0α expression revealed physical interactions between the GFP^+^ human photoreceptor precursors differentiated to M/L cones (characterised by OPN1LM/MW expression) and host bipolar cells in retinal wholemounts (**Fig. 2A’-A’’**). We also explored G0α expression in cross sections, which revealed intense expression associated with ON bipolar dendrites that appeared to contact GFP^+/^ OPN1LW MW cells in the INL (**Fig. 2B’-B’’, arrows**).

**Fig. 2.**
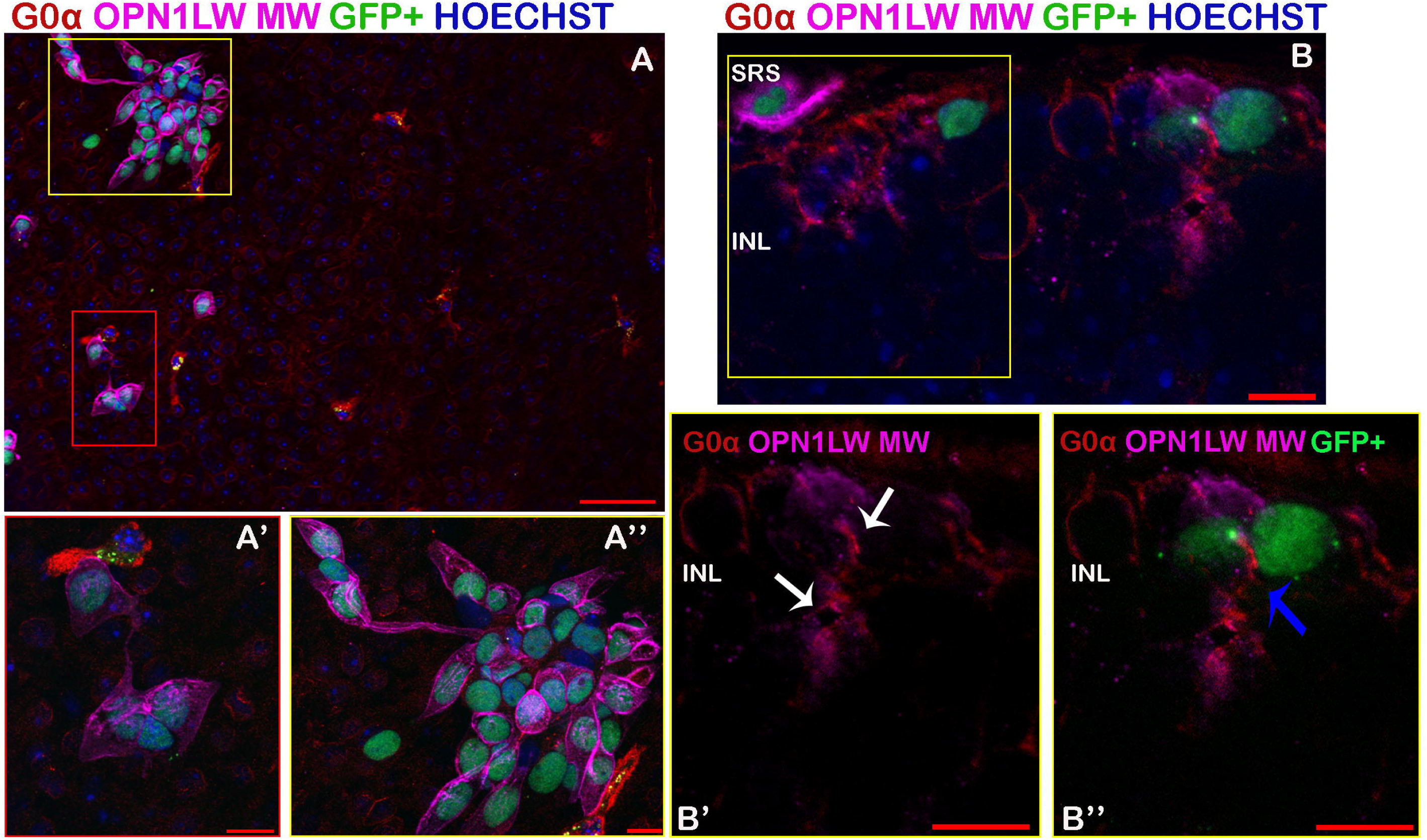
The transplanted CRX^+^ cells are found in close contact with the host bipolar cells. **(A)** Immunohistochemical analysis for G0α in red, opsin red\green (OPN1LW\MW) in pink and GFP^+^ cells in whole mount retina of *Pde6brd1* mice transplanted with CRX^+^ cells. (**A’-A’’**) Inset: high magnification images showing the GFP^+^-OPN1LW\MW expressing human donor cells found in close connection with the host cone bipolar cells. (**B**) The GFP^+^ and OPN1LW\MW immunostained cells were found in close apposition with the G0α host cells in the INL. Inset: (**B’**) high magnification images showing the anatomical interaction between the opsin red\green cells and the host cone bipolar cells (white arrows); (**B’’**) Representative image showing the colocalization between the GFP^+^ - OPN1LW\MW cells found in direct contact with G0α host cells (blue arrow). Scale bars, 50 µm (A, B); 10 µm (A’, A’’, B’ and B’’). Abbreviations: GFP, green fluorescent protein, INL, inner nuclear layer, SRS, subretinal space. Note: GFP in (A, A’, A’’, B and B’’) represents endogenous GFP expression.

In order to assess transplanted cell maturation and functional integration into host circuitry, we investigated restoration of visual function using two well-established visual behavioural tests, the light avoidance test (**Fig. 3A**) and the optomotor response test (**Fig. 3B**). As mice are nocturnal animals, they naturally tend to avoid bright environments (Misslin et al., 1989), a useful measure of visual function. Animals without functional photoreceptors therefore show less propensity for darkness. Behavioural testing was conducted after overnight dark adaptation using a white light source (∼1300 lux) in the illuminated half of the testing arena. Fifty percent of the transplanted mice (N=15) responded positively (**Movie S1**) and spent at least 5 out of 10 minutes (the total test duration) in the dark, with an average of 71.18% ± 14.21 (STD) of the test duration in the dark compartment (**Fig. 3C**). On the contrary, all the sham transplanted mice (N=10) spent more time in the bright area, resulting in 25.12% ± 0.84 (STD) of the total test duration in the dark compartment, reflecting the spontaneous exploratory behaviour of rodents in response to a novel environment (Bourin and Hascoët, 2003) (**Movie S2**).

**Fig. 3.**
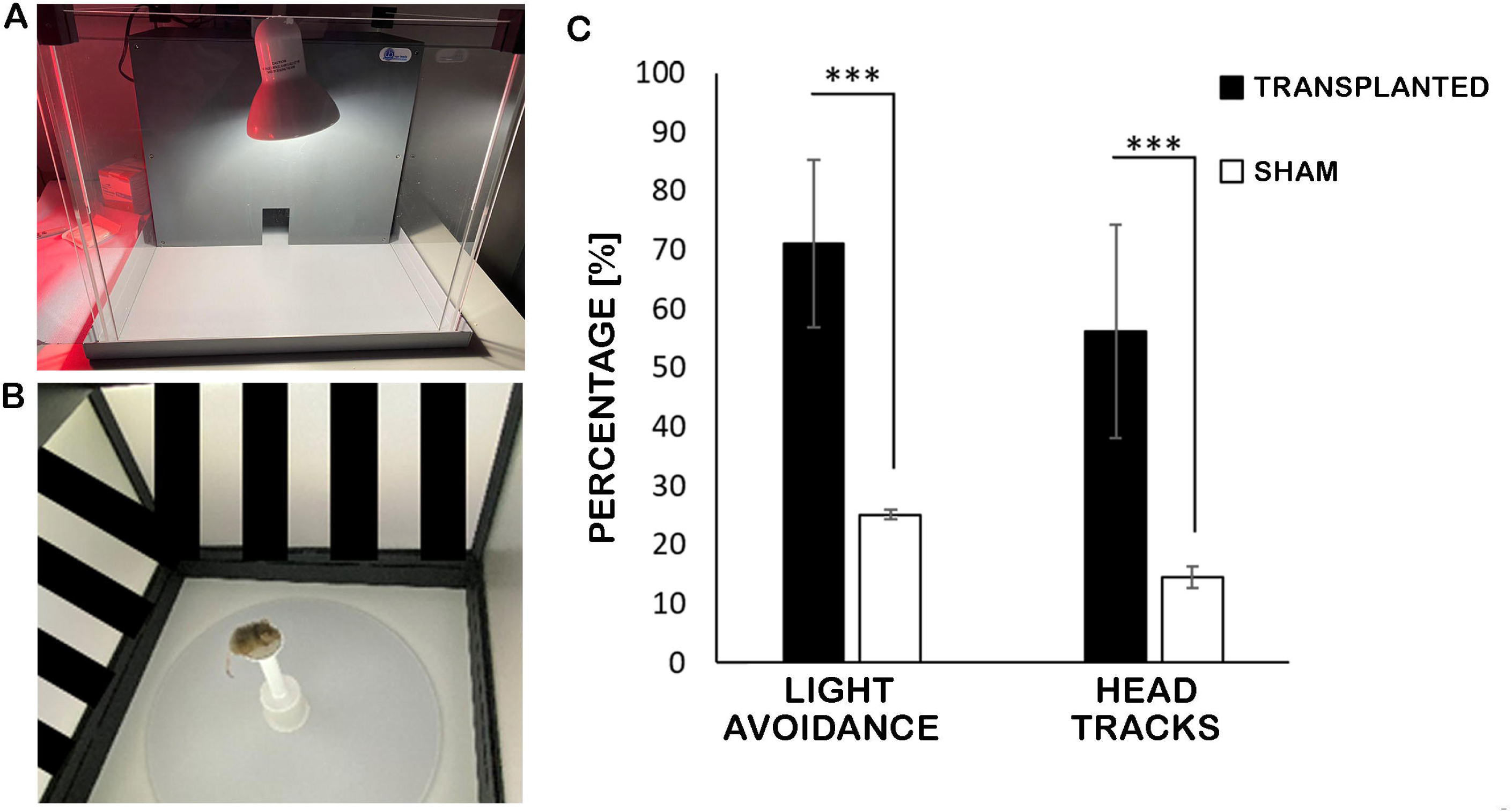
Light avoidance and Optomotor response. **(A)** Setup for the light avoidance test; (**B**) Photograph of the optomotor response test area; (C) Bar chart showing the significant differences (p< 0.001) in the light avoidance test and Optomotor response between the transplanted mice and the sham treated group. Data are presented as mean ± STD; statistical significance was assessed with Student’s t-test: ***P < .001.

Animals that exhibited light avoidance behaviour were tested for the optokinetic head tracking response to a rotating grating to find out whether they are able to discriminate motion and visual contrasts under photopic conditions. Tracking in each direction is independently driven by one eye (clockwise and anticlockwise by the left and by the right eye respectively) (Barnea-Cramer et al., 2016; Harvey et al., 1997). Since mice were transplanted bilaterally, responses to leftward versus rightward moving gratings can give us useful information about functionality in each eye and the software randomly decided the direction of the grating rotation. Transplanted mice exhibited significantly more head tracking (56% ± 18.08 (STD), N=15) than sham treated animals (14% ± 1.79 (STD) N=10) (**Fig. 3C**). As expected, low head-tracking behaviour was observed in sham transplanted mice, which can be attributed to the spontaneous and involuntary head movements.

### Light sensitivity is partially restored in CRX^+^ transplanted retinas

The mice that tested positive in the light-avoidance behavioural test were dark-adapted overnight and prepared for high-density large-scale MEA retinal recordings the next day. Rewiring of the neural retina in photoreceptor dystrophies results in permanent RGC oscillatory activity that masks light responses because of the poor signal-to-noise ratio. An example MEA raw data-trace of such spontaneous oscillations and bursts (5Hz high-pass filtering) is illustrated in in **Fig. 4A** (upper left trace). The underlying spurious spikes are still visible at 50Hz high-pass filtering when the slow oscillations are filtered out, masking potential light-driven responses (**Fig. 4A**, bottom left trace). Adding the gap junction blocker meclofenamic acid (MFA) dampens the oscillations and reduces the incidence of these spurious spikes (**Fig. 4A**, upper right trace), while preserving light responses (**Fig. 4C**) (Barrett et al., 2015, 2016). Moreover, addition of 40 µM MFA has no effect on the spike waveform shape, which is crucial for accurate isolation of single units (**Fig. 4B**). To illustrate the signal-to-noise improvement with MFA, light driven activity of the RGC from Panel 4B is shown without and with MFA in Panel 4C. The cell was exposed to 50 repetitions of the same full field stimuli series (trials on the y-axis) and the spike rasters and post-stimulus time histograms for that particular unit are compared without (left) and with (right) 40 µM MFA application. Before adding the drug, it is barely possible to see that this cell responds to the onset of the light (**Fig. 4C**, left). With MFA, the baseline-firing rate significantly drops, which vastly improves the signal-to-noise ratio, revealing clear ON responses (**Fig. 4C**, right).

**Fig. 4:**
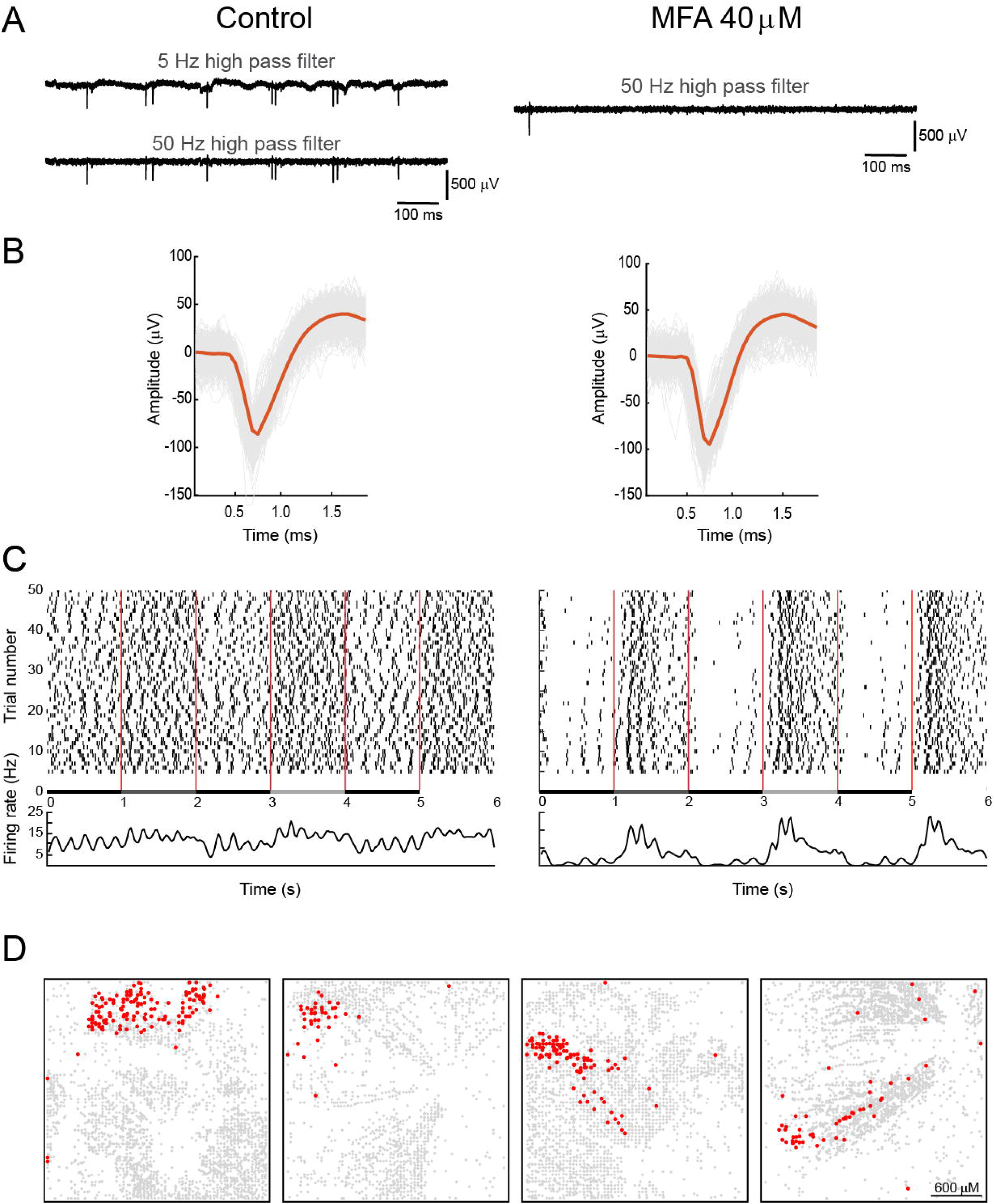
Dampening RGC oscillatory activity with MFA enhances the expression of light responses. **(A)** Left: Raw data traces (from one electrode of the 4096 channels MEA) from a *Pde6brd1* retina in control conditions filtered with a 5 (top) and 50 (bottom) Hz high-pass filter. Right: Raw data trace recorded (50 Hz high-pass) from a *Pde6brd1* retina in 40 µM meclofenamic acid (MFA). (**B**) Extracted spike waveforms (grey plots) from one isolated RGC unit (the same unit as shown in C) in control conditions (left) and in the presence of 40 µM MFA (right). The mean spike waveform is plotted in red. (**C**) Spike raster (top) and post-stimulus histograms (PSTH, bottom) for the same RGC unit in control (left) and 40 µM MFA (right). Spike raster: Spike times of 50 repetitions with the same stimulus (1-second contrast steps, indicated between spike raster and PSTH) are plotted as small vertical ticks. The average rate (spikes/s) of these repetitions is represented on the y-axis of the PSTH (bin size = 25 ms). (**D**) The x- and y-coordinates of all spike location centres, obtained with the spike sorting package Herdingspikes2, are plotted in grey in a 64 × 64 grid (the electrode layout of the high density MEA). The units that were identified as light responsive are plotted in red.

We screened six mice (12 retinas) for clear visible light responses (with 40 µM MFA) to full field stimuli with increasing contrasts and square wave gratings with different bar widths and directions. We identified seven retinas (from five different mice) with RGCs showing clear responses to light (full field flashes, motion or both). To visualise the spatial location of these light-responsive RGCs with respect to the recording area, we plotted the x, y coordinates of all active units on the MEA in four representative explanted retinas (**Fig. 4D**). The red dots represent the light-responding units, showing that these responsive cells always cluster in a restricted area (presumably corresponding to the injection site in the temporal retina). The grey dots show all the other RGCs with residual spontaneous activity but no light responses. The average sample size per experiment for recorded RGCs from these seven retinas was 2910±928 (STD). On average, 45±36 RGCs or 1.73±1.35 % of the entire recorded population exhibiting neural activity also responded to light.

Overall, we observed mainly “conventional” ON, OFF and ON-OFF responses. To illustrate responses to full field flashes (**Fig. 5**, left panels) and to different square wave gratings (**Fig 5**, right panels), we selected one representative cell for each of these conventional types. Many of the cells exhibited contrast sensitivity, suggesting that some of the basic retinal circuitry is preserved despite neural remodelling (for example, **Fig. 5A** with clear increase in responsiveness for gradually brighter stimuli). Accordingly, many of these cells showed gradually increasing response suppression with an increase in the grating spatial frequency (**Fig. 5B, D, F**) or, alternatively, had their maximum activity at a specific bar width (**Fig. 5H**). A few cells showed significantly higher activity towards a given direction (**Fig. 5H**, purple), but often only for a specific bar width (**Fig. 5B**, 500 µm, green).

**Fig. 5:**
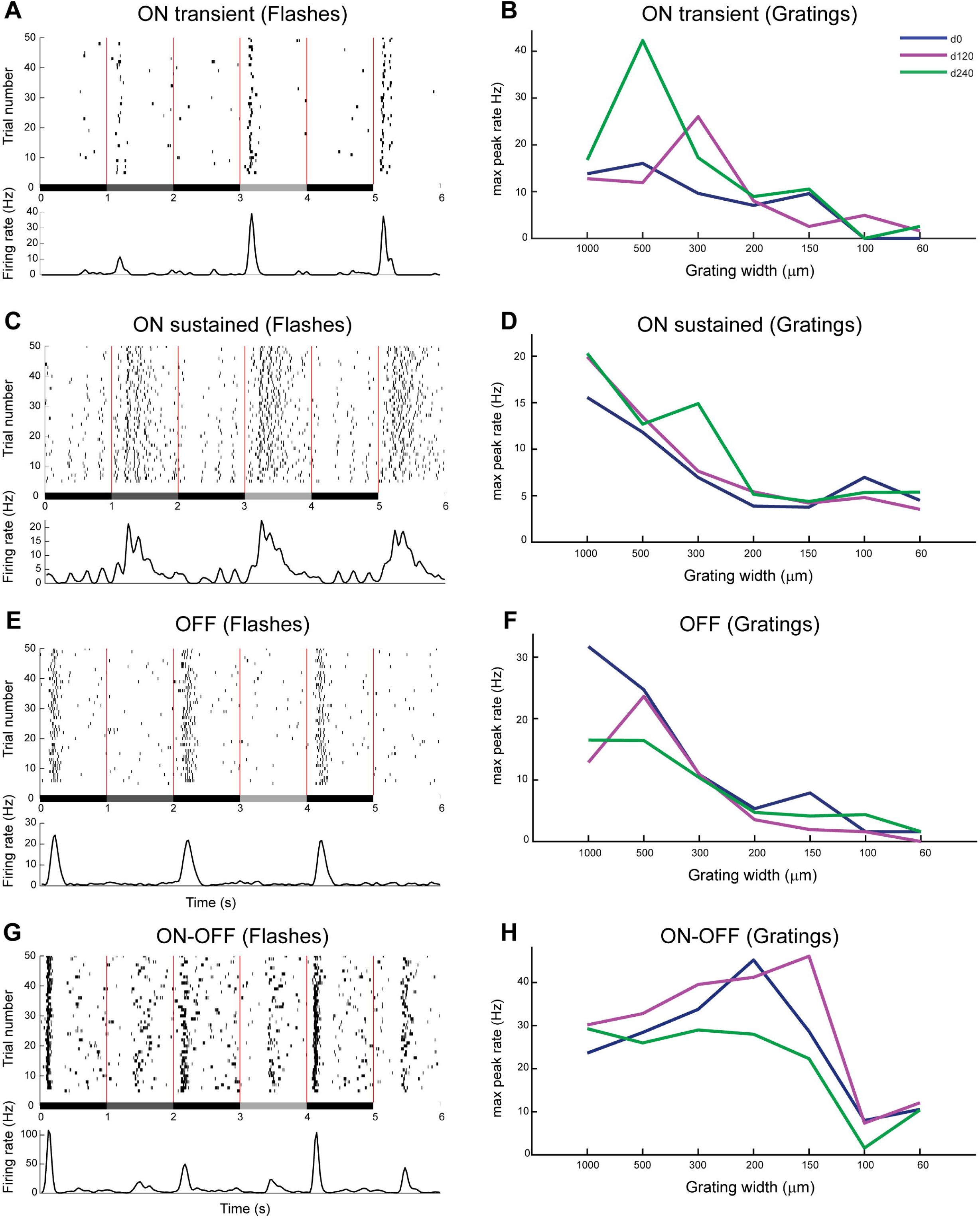
Conventional light responses. Example spike rasters and PSTHs from selected RGCs to 1-second contrast steps (flashes) of (**A**) ON transient, (**C**) ON sustained, (**E**) OFF and (**G**) ON-OFF responses. The maximum responses to square wave gratings with different bar widths (x-axis) and directions (d0 = blue, d120 = magenta, d240 = green) are plotted for the same units as in A, C, E and G (**B** = ON transient, **D** = ON sustained, **F** = OFF, **H** = ON-OFF).

However, we also observed substantial numbers of responses that are never encountered in healthy, normal retinas (**Fig. 6**). These unusual responses can be categorised into four main groups. Three of these groups share OFF excitatory responses and pronounced activity suppression induced at the light onset (ON suppressed or suppressed by light (SBL)). SBL OFF trans cells (**Fig. 6A**) showed a transient and robust response to light offset, SBL OFF sust cells (**Fig. 6B**) showed a more sustained response to light offset and SBL OFF delayed cells exhibited a delayed response to light offset (**Fig. 6C**). The fourth group exhibited ON sustained responses with dark-suppressed activity (SBD ON sust, **Fig. 6D**). We also found a few cells with faint responses to light, only visible during moving gratings, but not during static flashes (**Fig. 7**, only motion). Control experiments with sham injected *Pde6brd1* retinas did not show light responses at all. The relative incidence of the different functional groups we recorded from for all retinas is summarised in **Fig. 7** (details given in legend). Most of the responses were conventional (63.04%) while the unusual response types made 22.05% and the DS/only motion types 14.91% of all light responses.

**Fig. 6:**
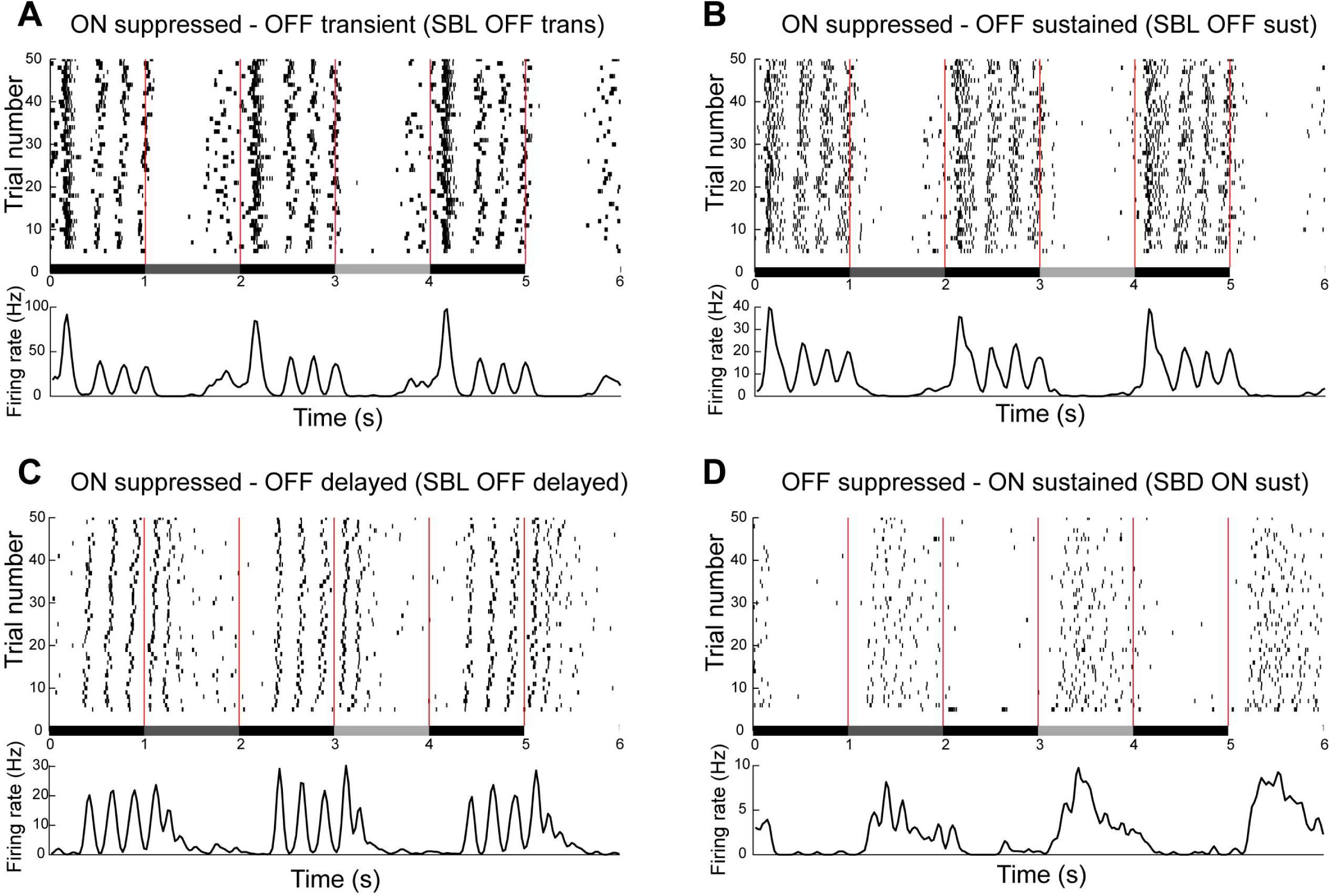
Unconventional light responses. Example spike rasters and PSTHs from selected RGCs to 1-second contrast steps (flashes) of (**A**) ON suppressed – OFF transient (SBL OFF trans), (**B**) ON suppressed – OFF sustained (SBL OFF sust), (**C**) ON suppressed – OFF delayed (SBL OFF delayed) and (**D**) OFF suppressed – ON sustained (SBD ON sust).

**Fig. 7:**
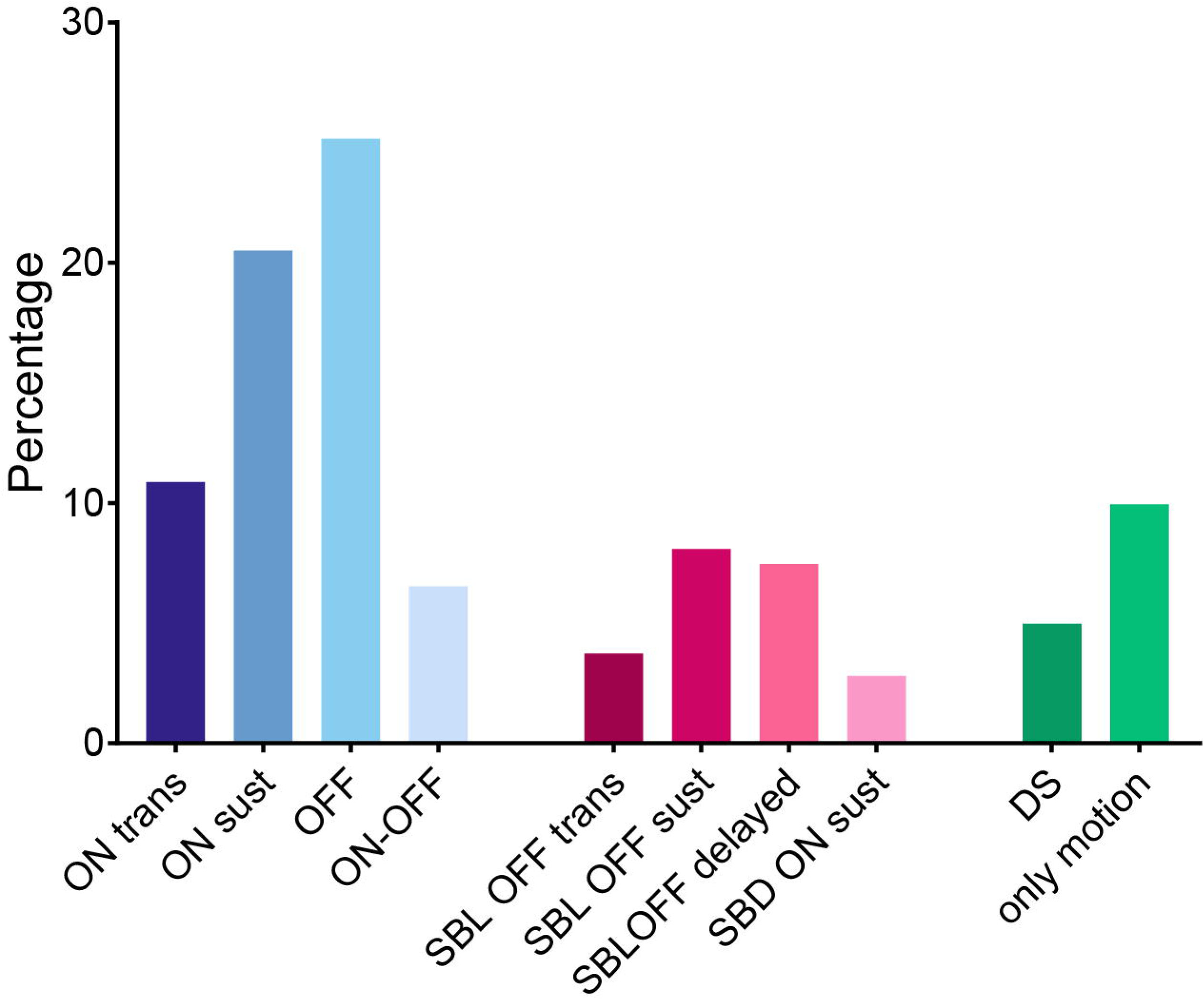
Summary of light responses grouped into four conventional response clusters. (ON trans, ON sust, OFF, ON-OFF), four unconventional response types (SBL OFF trans, SBL OFF sust, SBL OFF delayed, SBD ON sust) **and DS/only motion**. The response types were counted from the 7 light responsive retinas and values were calculated as percentage from all responding cells. In detail: ON trans (10.87%), ON sust (20.5%), OFF (25.16%), ON-OFF (6.52%) SBL OFF trans (3.73%), SBL OFF sust (8.08%), SBL OFF delayed (7.45%), SBD ON sust (2.8%), DS (4.97%) and only motion (9.94%).

To further characterise the nature of the conventional and unusual responses, we conducted additional experiments with L-AP4, an agonist for group III metabotropic glutamate receptors and with DNQX, an antagonist for AMPA/kainate ionotropic glutamate receptors. The drugs were applied in succession, starting with 20 µM L-AP4 to hyperpolarize ON bipolar cells, followed by 20 µM L-AP4 + 20 µM DNQX to additionally block AMPA/kainate ionotropic receptors in OFF bipolar cells and other retinal neurons. DNQX indeed abolished all remaining light responses (data not shown). **Fig. 8** compares the light responses in control conditions versus in the presence of 20 µM L-AP4. As expected, ON transient RGCs (**Fig. 8A**), did not show any light responses in the presence of L-AP4, even though there was an increase in spontaneous activity levels (**Fig. 8B**). OFF responses were still present in L-AP4 (**Fig. 8D**) but not as conspicuously as in control conditions (**Fig. 8C**). SBL OFF transient (**Fig. 9E, F**) and sustained (**Fig. 8G, H**) responses behaved in a similar pattern - the initial OFF response decreased in L-AP4. Further, the delayed OFF responses, which normally follow the initial OFF responses (**Figure 8E, G**), were less sensitive to the drug (**Fig. 8F, H**).

**Fig. 8:**
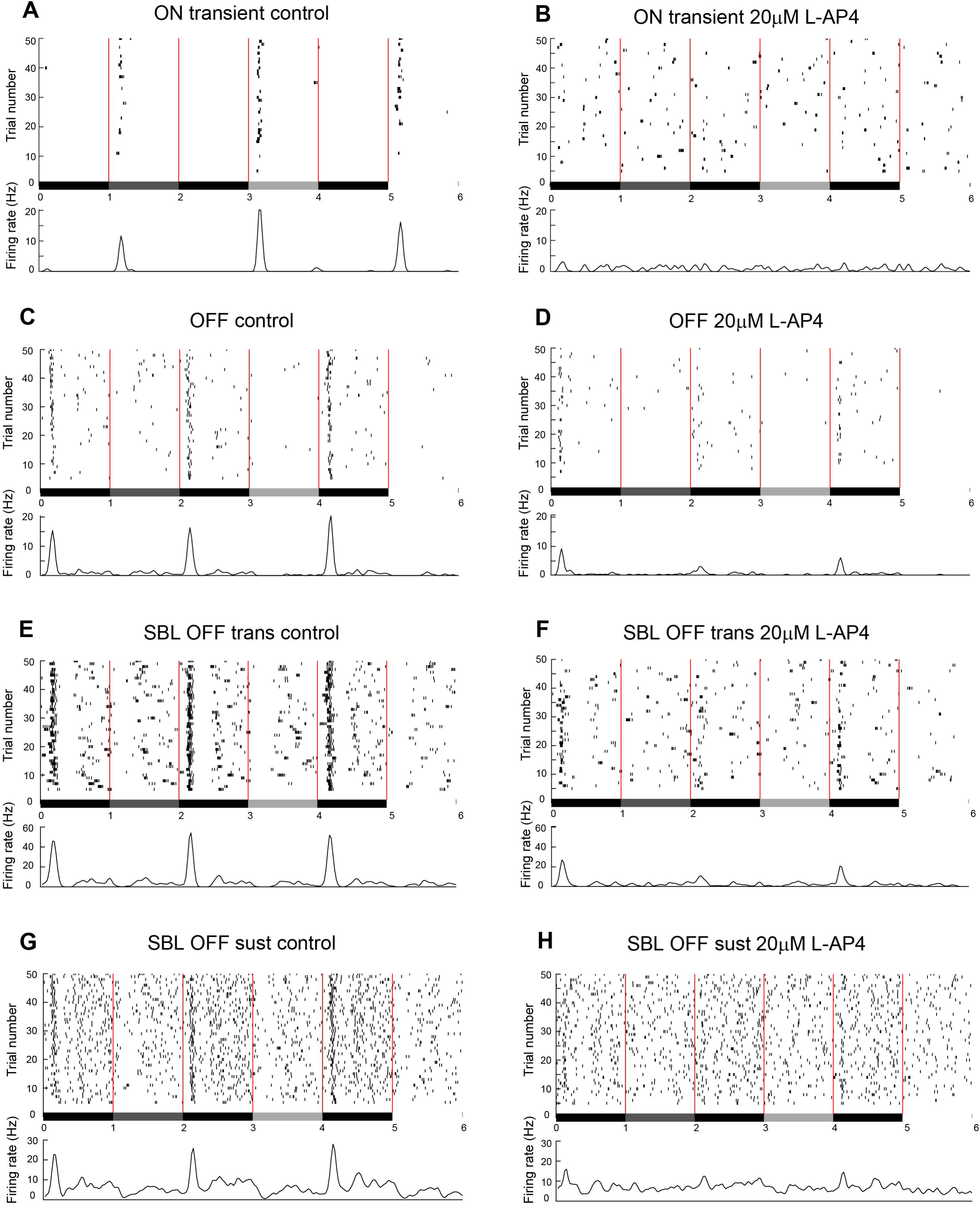
Responses in control and during activation of mGLUR6 receptors with L-AP4. Example spike rasters and PSTHs from selected RGCs to 1-second contrast steps (flashes) in control (**A, C**, **E, G**) and 20 µM L-AP4 (**B, D, F, H**). (**A, B**) ON transient response; (**C, D**) OFF response; (**E**, **F**) Supressed by light (SBL) OFF transient response; (**G, H**) SBL OFF sustained response.

In summary, our data show that transplanted human photoreceptor cells are able to elicit conventional light responses from host RGCs in the majority of cases. Unconventional response types were also present at a lower incidence, which we attribute to rewiring of the inner retina that occurs during early degeneration stages, resulting in abnormal connectivity.

## DISCUSSION

We have developed robust protocols for generation of photoreceptors within pluripotent stem cell derived retinal organoids (Dorgau et al., 2019; Mellough et al., 2015; Zerti et al., 2020) and have shown that these are able to engraft into animal models of retinal degeneration (Collin et al., 2019). Notwithstanding, there are several questions that remain to be investigated with regard to establishment of a layer of light-sensitive cells at such density that allows integration of transplanted photoreceptors within the host retinas and enables visual improvement. In this study, we investigated the ability of transplanted hESC-derived cone precursors to elicit light responses in adult RGCs in the *Pde6brd1* mouse, characterised by advanced retinal degeneration. Our approach was to measure light-driven activity from the RGC layer using high-density, large-scale MEAs that allow simultaneously recording from hundreds to thousands of RGCs over a large area. Our data indicate that 80% of subretinally transplanted mice show engraftment of hESC-derived CRX-GFP^+^ cone precursors. Up to 1.5% of cells integrated into the putative host ONL, differentiated into cones and were able to form synapses with host bipolar cells. This integration efficiency is similar to that found in wild-type or other models of cone degeneration transplanted with cone photoreceptors, as recently reported by other groups (Santos-Ferreira et al., 2015; Smiley et al., 2016).

Of the transplanted mice, 50% were able to perform the behavioural tests and 33% showed light sensitivity in both eyes, demonstrating restoration of light responses in *Pde6brd1* mice. Normally, both rods and cones degenerate by the end of the first postnatal month in this mouse model of retinitis pigmentosa, resulting in complete lack of light responsiveness in all retinal neurons. Four weeks following transplantation, we were able to consistently record responses to various light stimuli, including responses to different light intensities and to motion. Rather than being scattered all over the retina, the responsive cells were always clustered in one area, presumably reflecting the single cone progenitors’ injection site. We are confident that these temporally precise, stimulus-locked responses do not arise from remaining intrinsically photosensitive RGCs (ipRGCs) as the latter have very poor temporal fidelity (Procyk et al., 2015) in photoreceptor dystrophies. Moreover, ipRGCs are scattered all over the RGC layer, whilst we always saw responses only in a restricted area.

In additional support, ON responses disappear in the presence of the mGluR6 agonist L-AP4 and ON and OFF responses disappear in the presence of L-AP4 + DNQX (AMPA/kainate receptor antagonist), indicating that these responses are exclusively driven by glutamatergic neurotransmission, and not by ipRGCs. It is also unlikely that the responses we recorded originate from residual S and M cones reported in previous studies (Lin et al., 2009; Narayan et al., 2019). Indeed, after week 12, residual S cones are restricted to the far periphery of the ventral retina, while remaining M cones are located in the far periphery of the dorsal retina (Lin et al., 2009); furthermore, these remaining cones are morphologically abnormal and lack inner and outer segments (Barnea-Cramer et al., 2016). Though our responsive RGCs cluster in spatially restricted areas, these areas always stretch from the periphery towards the centre, lending support to the fact that they do not reflect surviving cones in the periphery. Finally, none of the sham-injected retinas exhibited any clear light responses.

Photoreceptor transplantation approaches to rescue visual function has made a significant leap in recent years (Gasparini et al., 2019). For example, Santos-Ferreira et al. (Santos-Ferreira et al., 2015), performed cone-like cell transplantation in a cone degeneration mouse model and recorded ON and OFF, and to some extent ON-OFF RGC responses in day-light conditions. Recently, Mandai et al. (Mandai et al., 2017) transplanted mouse iPSC-derived retinal grafts at end-stage degeneration in *Pde6brd1* mice (7-9 weeks) and recorded simple, mainly transient ON responses to light in RGCs 2-4 months after transplantation. Both studies found mainly ON and OFF light responses whereas our results show a larger variety of responses including clear transient and sustained ON, OFF and ON-OFF responses shortly after transplantation. It is astonishing, but at the same time not surprising, that despite the profound morphological, biochemical and physiological impact caused by retinal remodelling, a large set of conventional response types are still present. It is known that the surviving retinal circuitry in photoreceptor-degenerated retinas are preserved, at least up to a certain stage of degeneration, allowing for the generation of conventional ON, OFF and ON-OFF responses following subretinal electrical stimulation (*Pde6brd1* retinae < P56 (Haq et al., 2018; Stutzki et al., 2016)). Here, however, we report photoreceptor-driven ON, OFF and ON-OFF responses in CRX+ transplanted *Pde6brd1* mice older than 12 weeks, suggesting that retinal circuitry is somehow preserved for longer than previously assumed.

Importantly, we show for the first time unconventional responses, which potentially result from novel, aberrant neural wiring that develops during remodelling of the neural retina. The ON and OFF pathways are differently affected during the progressive retinal remodelling (Stasheff, 2008) and Jones et al. (Jones et al., 2003) describe that some cells are less affected, showing signs of rewiring and de *novo* synapse formation. This suggests a certain degree of plasticity and it is likely that transplanted cones induce the formation of functional contacts with host neurons, for example bipolar cells, or perhaps even contact RGC dendrites directly, which could explain the strong inhibitory responses to light we have observed in a substantial number of cells. Indeed, our immunohistochemistry data suggests that the GFP labelled transplanted cone precursors successfully engraft and make potential contacts with bipolar cells.

In summary, our study shows that hESC-derived cone precursors engraft and elicit both conventional and unconventional light responses in mice with advanced retinal degeneration, despite a relatively low number of engrafted cells. This demonstrates that hESC-derived cones provide a useful source of cells for transplantation into demised retinas; however to progress this to human clinical trials, further work is needed to improve the number of engrafted cells and slow down remodelling of the neural retina to achieve clinically useful vision improvement in patients with degenerative retinal diseases.

## MATERIALS AND METHODS

### Human pluripotent stem cell culture and differentiation

The human embryonic stem cell (hESC) line harbouring the GPF reporter at the 3’UTR of CRX locus was expanded in mTeSR™1 (StemCell Technologies) at 37°C and 5% CO2 on 6 well plates pre-coated with Low Growth Factor Matrigel (Corning). Differentiation to retinal organoids was performed as described in Collin et al., 2019. Retinal organoids were collected on day 90, dissociated to single cells and subjected to flow activated cell sorting to enable enrichment of CRX-GFP^+^ for transplantation.

### Subretinal transplants

#### Ethics statement

All experimental work performed in this study was in accordance with the United Kingdom Animals (Scientific Procedures) Act 1986 and carried out in accordance with protocols approved by the Animal Welfare and Ethics Committee of Newcastle University. All efforts were made to minimize the number and the suffering of animals used in these experiments.

#### Animals

*C3H/HeNHsd*-*Pde6brd1* mice without the *Gpr179* mutation were kindly gifted from Professor Robin Ali’s lab (UCL London). The mice were housed in the Animal facility at Newcastle University on a standard 12-hour light/dark cycle at the same light levels throughout the experimental period. Animals were kept in ventilated cages with food and water *ad libitum*.

#### Immune Suppression

To prevent immune-rejection of the transplanted human cells, daily subcutaneous injections of Cyclosporine A (50 mg/kg/day) were administered to the recipient animals starting at 1 day before the transplantation and maintained throughout the experiment.

#### Surgery and transplantation

Surgery was performed, in male and female mice aged ∼10-12 weeks, under direct ophthalmoscopy through an operating microscope, as previously described (Collin et al., 2019). 100, 000 CRX^+^ cells in a volume of 1 µl volume were injected in each eye.

### Behavioural Tests

#### Light Avoidance tests

A light-chamber/dark chamber apparatus (Ugo Basile, Ltd) was used to assess the preference for light or dark compartment in male and female mice that had received bilateral CRX-GFP^+^ cell transplants (N=28) or bilateral sham injection (N=5) and control *Pde6brd1* mice that had received no cell transplants (N=5). Mice were tested in a custom-made area measuring 26 × 26 × 26 cm containing equally sized dark and light chambers connected by a 4-× 5-cm aperture at midpoint. The mice were dark-adapted >12 h before testing. Testing was conducted in a dark room, and the light chamber was lit by a custom LED array suspended above the chamber emitting white light at 1300 lux. Both compartments had removable tops to allow thorough cleaning with 70% ethanol before each test. Both pupils were dilated with one drop of 1% atropine instilled ∼10 minutes before testing. Under dim red light, each mouse was placed in the light compartment with the light turned off, ensuring minimal tail restraint, and gently placed in the middle of the chamber to be lit (front half), facing away from the connecting aperture. The experimenter was masked to treatment group. The chambers were closed, LED light was turned on, and video recording commenced. Each trial lasted 10 minutes, and all mice were test-naive (one trial per mouse). A mouse was deemed to have entered a chamber when four paws had crossed into that compartment. Mice were assayed for time spent in the lit chamber and data were calculated manually by viewing the video recording.

#### Optomotor Response

We measured visual contrast sensitivity of mice by observing their optomotor responses to moving sinewave gratings randomly by the OptoMotry^™^ Software (Prusky et al., 2004). Mice reflexively respond to rotating vertical gratings by moving their head in the direction of grating rotation. The protocol used yields independent measures of the acuities of right and left eyes based on the unequal sensitivities of the two eyes to pattern rotation: right and left eyes are most sensitive to counter-clockwise and clockwise rotations, respectively. A virtual-reality chamber is created with four 17-inch computer monitors facing into a square. As shown in **Fig. 3B**, a virtual cylinder, covered with a vertical sine-wave grating, was drawn and projected onto the monitors using the Optomotry software on an Apple Power Macintosh computer. The animal was placed on a platform in the centre of the square and a video camera, situated above the animal, provided real-time video feedback on another computer screen. From the animal perspective, the monitors appeared as large windows through which it viewed the rotating cylinder. A red crosshair in the video frame indicated the centre of the cylinder rotation. Each tested mouse was placed on the platform, the lid of the box was closed, and the animal was allowed to move freely. As the mouse moved about the platform, the experimenter followed the mouse’s head with a crosshair superimposed on the video image. A trial started when the experimenter cantered the virtual drum on the head; a drifting (12 deg/s) grating then appeared. When a grating perceptible to the mouse was projected on the cylinder wall and the cylinder was rotated, the mouse normally stopped body movements and began to track the grating with reflexive head movements in concert with the rotation. The experimenter judged whether the mouse made slow tracking movements to follow the drift grating. Large repositioning and grooming movements were ignored, and the trial was restarted if tracking movements were not clearly seen. All animals were habituated before the outset of testing with handling and by placing them on the platform for a few minutes at a time. The mice were generally tested for 5-20 minutes during the first few hours of their daylight cycle. Our typical procedure was to have an experimenter make a judgment on each trial about whether the animal tracked. Although this has proved highly efficient and enables thresholds to be measured quickly, it opens the possibility of experimenter bias affecting the results. To check whether this was a real problem, for one experiment we altered the system so that the experimenter could not control the stimulus direction. Drum rotation was random from trial to trial, and the experimenter had to make a forced-choice decision between clockwise and counter-clockwise rotation. After placing the animal on the platform and closing the lid, the experimenter waited until the animal stopped moving, at which time the gray uniform background was replaced with a 0.05 cycle\degrees spatial-frequency sine wave grating (100% contrast) under photopic condition (60 cds\m^2^) and moving in one direction. In order to assess visual acuity, gratings had a constant contrast of 100%. Using a staircase paradigm, the program converges to measures of the contrast sensitivity of both eyes defined as the spatial frequency or % contrast yielding ≥ 70 % correct observer responses (Umino et al., 2008). The animal was assessed for tracking behaviour for a few seconds, and then the gray stimulus was restored. This procedure was repeated until unambiguous tracking was observed. The short testing epochs reduced the possibility of the mouse’s adapting to the stimulus and established that each animal was capable of tracking when a salient stimulus was present, and the testing with a fixed spatial frequency and contrast grating enabled each mouse’s optomotor response to be properly evaluated. Occasionally, during testing, sudden squeaking noises, or taps on the lid were interspersed with the grating presentations to induce the animal to stop moving, which facilitated more rapid testing.

#### Histology and Immunohistochemistry

Animals were sacrificed by cervical dislocation. Eyes were quickly removed and dissected in 4% paraformaldehyde (PFA) in PBS. Eye cups were prepared and fixed for 1 hour at 4°C, incubated overnight in 30% sucrose solution, and embedded in OCT. Cryosections were cut at 20 µm thick and all sections were collected onto Superfrost Plus slides and store at -20C before the staining. Retinal section or whole mount retinas were hydrated with PBS for 30 minutes and incubated with blocking solution containing 10% goat serum and 0.3 % Triton-X-100 for 1 hour at RT. Slides were incubated with the appropriate primary antibody overnight 4°C (**Table S1**). After rinsing with PBS, sections were incubated with the secondary antibody for 2 hours at RT, rinsed and counter-stained with Hoechst. Alexa Fluor 546 and 647 secondary antibodies (Invitrogen-Molecular Probes) were used at a 1:1000 dilution. Negative controls were carried out by omitting the primary antibody.

#### Image acquisition and processing

Retinal sections and whole mounts retina were viewed on a Zeiss Axio ImagerZ2 equipped with Apotome 2 and Zen 2012 blue software (Zeiss, Germany). Objectives lens used were EC Plan Neofluar 20x/0.5 Ph2 and EC Plan Apochromat 63x/1.4 Ph3. Series of XZ optical sections (<1 µm thick) were taken at 1.0 µm steps throughout the depth of the section. Final images are presented as a maximum projection and adjusted for brightness and contrast in Adobe Photoshop CS6 (Adobe Systems).

### Electrophysiology

#### Retina Preparation for electrophysiological recordings

All experimental procedures were approved by the ethics committee at Newcastle University and carried out following the guidelines of the UK Home Office, under control of the Animals (Scientific Procedures) Act 1986. Male and female *Pde6brd1* mice aged between postnatal weeks 14-18, which passed the light avoidance and optomotor tests, were used for *in vitro* recordings from the RGC layer. Mice were dark-adapted overnight and killed by cervical dislocation. Eyes were enucleated, and following removal of the cornea, lens, and vitreous body, they were placed in artificial cerebrospinal fluid (aCSF) containing the following (in mM): 118 NaCl, 25 NaHCO_3_, 1 NaH2 PO_4_, 3 KCl, 1 MgCl_2_, 2 CaCl_2_, 10 glucose, and 0.5 l-Glutamine, equilibrated with 95% O_2_ and 5% CO_2_. The dorsal orientation was marked after enucleation. The retina was isolated from the eye cup and flattened with the temporal area, the transplantation side, covering the active recording area of the MEA. Retinas were allowed to settle for at least 2 hours before recording. All procedures were performed in dim red light and the room was maintained in darkness throughout the experiment.

#### High-density large-scale MEA recordings

Recordings were performed on the BioCamX platform with HD-MEA Arena chips (3Brain GmbH, Lanquart, Switzerland), integrating 4096 square microelectrodes in a 2.67 × 2.67 mm area. The platform records at a sampling rate of ∼18 kHz/electrode when using the full 64×64 array and recordings were stored in an hdf5-based data format either with a 5 or 50 Hz high-pass filter using 3Brain’s BrainWaveX software application. Spikes were extracted from the raw traces using a quantile-based event detection (Muthmann et al., 2015). Single-unit spikes were sorted using Herdingspikes2 (https://github.com/mhhennig/HS2), an automated spike sorting method for dense, large-scale recordings (Hilgen et al., 2017).

Light stimuli were projected as described previously (Barrett et al., 2016). A full field stimulus with increasing contrasts (1-second contrast steps, max irradiance 217 µW/cm^2^, 1 second (s) black, followed by 1 s 33% grey, 1 s black, 1 s 66% grey, 1 s black and 1 s white) was used to detect potential basic light-driven ON-OFF responses. We also used a square wave grating with different bar widths (1000, 500, 300, 200, 150, 100 and 60 µM) presented at three different directions (0, 120 and 240 degrees) to unmask responses to motion and to test whether some RGCs were sensitive to the direction of motion (direction selectivity). The light intensity used in these experiments was set to high photopic (bright day light) conditions to ensure predominant cone activation. Bath applications with 40 µM MFA (Sigma Aldrich, ST. Louis, MO), 20 µM L-AP4 (Sigma Aldrich, ST. Louis, MO) and 20 µM DNQX (Tocris, Bristol, UK) were applied for 30 minutes before recording. Statistical significance and firing rate analyses were evaluated by using Prism (GraphPad, CA) and MATLAB (Mathworks, MA). Briefly, post-stimulus time histograms were calculated using Matlab’s histogram function. Activity plots were generated by using the scatterplot function and the x,y coordinates (provided by the spike sorter Herdingspikes2) of light responsive (red) and non-responsive (gray) cells (**Fig. 4D**).

### Statistical Analyses

All statistical tests were performed using Prism (GraphPad, USA). Statistical significance was tested using two-tailed Student’s t-test. * = p-value < 0.05, ** = p-value < 0.01, *** = p-value <0.001.

## Acknowledgments

We are grateful to ERC (#614620), RP Fighting Blindness (#GR593), Leverhulme Trust (RPG-2016-315), and BBSRC (BB/P018440/1) for funding this work. We would also like to thank Prof. Robin Ali for the kind donation of *Pde6brd1* mice and Dr. Kathy Murphy at the Newcastle Comparative Biology Centre for fruitful discussions on animal surgery and welfare.

## Author Contributions

D.Z. and G.H. experimental design, performed research, data collection and analysis, figure preparation and manuscript writing;

B.H., J.C. and L.A. performed research and data collection;

M.A. donated reagents, performed training in subretinal transplants and visual tests;

E.S. and M.L. study design, data analysis, figure preparation, manuscript writing and fund raising. All authors approved the final version of the manuscript.

## Table

**Table 1:** List of the antibodies used for the IHC.

## Supplemental Information

**Movie S1: Light-avoidance test behaviour after photoreceptor cell transplantation**. This *Pde6brd1* mouse had a positive result.

**Movie S2: Light-avoidance test behaviour after photoreceptor cell transplantation**. This *Pde6brd1* mouse had a negative result.

