## Supplementary material for "Transplanted pluripotent stem cell-derived photoreceptor precursors elicit conventional and unusual light responses in mice with advanced retinal degeneration": Table s1

**Table 1.** List of the Antibodies used for the IHC.

| **Antibody** | **Host** | **Dilution** | **Supplier, Cat. No** |
| --- | --- | --- | --- |
| Anti-Recoverin | Rabbit | 1:1000 | Millipore, AB5585 |
| Anti-opsin Blue (OPN1SW) | Rabbit | 1:200 | Millipore, AB5407 |
| Anti-opsin Red/Green (OPN1LW/MW) | Rabbit | 1:200 | Millipore, AB5405 |
| Anti-G Protein Goα | Mouse | 1:500 | Millipore, MAB3073 |
